## Supplementary material for "ATF3 Preserves Skeletal Muscle Stem Cell Quiescence by Preventing Precocious Activation": Suppl

**Inventory of Supplemental Information**

1. **Supplemental Figures**

Figure S1. ATF3 is rapidly and transiently induced during early SC activation.

Figure S2. Short-term *Atf3* deletion accelerates acute injury-induced muscle regeneration.

Figure S3. *Atf3* deletion provokes premature SC activation and pseudo-regeneration in homeostatic muscle.

Figure S4. Long-term *Atf3* deficiency depletes SC pool and impairs muscle regeneration.

Figure S5. ATF3 deletion induces SC activation during voluntary and resistance exercises.

Figure S6. ATF3 regulates *H2B* gene expression and nucleosome patterning.

Figure S7. Screening of other functional AP-1 family members in SCs and regeneration.

2. **Supplemental Tables**

Table S1. Transcriptomic profiling in *Atf3* iKO SCs.

Table S2. Genome-wide profiling of ATF3 binding.

Table S3. Transcriptomic profiling in *Atf3* cKO SCs.

Table S4. CUT&RUN analysis of H2B binding in *Atf3* iKO SCs.

Table S5. Information of oligonucleotides and primers used in the study.

3. **Supplementary figure legends**

**Supplementary Figure 1. ATF3 is rapidly and transiently induced during early SC activation. A** qRT-PCR detection of *Atf4*, *Fos*, *Fosb* and *Junb* in QSC, FISC and ASC-24h from muscles of Tg: *Pax7-nGFP* mice. **B** Immunofluorescence staining of ATF4, FOS, FOSB or JUNB and PAX7 proteins on QSCs and FISCs. Scale bar: 50μm. All the bar graphs are presented as mean ± SD. ∗∗p < 0.01, ∗∗∗p < 0.001.

**Supplementary Figure 2. Short-term *Atf3* deletion accelerates acute injury-induced muscle regeneration. A&B** IF staining for Pax7 and ATF3 on (A) FISCs or (B) single myofibers from Ctrl or *Atf3* iKO mice. Scale bar: 50μm. **C** Left: representative images of TA muscles from Ctrl or *Atf3* iKO mice 30 days post the 1^st^ round injury. Right: Quantification of the above TA muscle weight; n = 3 mice per group. **D** Left: H&E staining of the above TA muscles. Scale bar: 50 μm. Right: Quantification of the average CSAs of newly formed fibers; n = 3 mice per group. **E** Left: representative images of TA muscles from Ctrl or *Atf3* iKO mice 30 days post the 3^rd^ round injury. Right: Quantification of the above TA muscle weight; n = 3 mice per group. **F** Left: H&E staining of the above TA muscles. Scale bar: 50 μm. Right: Quantification of the average CSAs of newly formed fibers; n = 3 mice per group. All the bar graphs are presented as mean ± SD. ∗p < 0.05, ∗∗p < 0.01, ∗∗∗p < 0.001.

**Supplementary Figure 3. *Atf3* deletion provokes SC premature activation and pseudo-regeneration in homeostatic muscle. A** Left: IF staining of Myod (green) and MyoG (red) on SCs isolated from Ctrl or iKO mice and cultured for 72h. Scale bar: 50 μm. Right: Quantification of the percentage of Myod+/MyoG+ SCs on single myofibers; n = 3 mice per group. All the bar graphs are presented as mean ± SD. ∗p < 0.05.

**Supplementary Figure 4. Long-term *Atf3* deficiency depletes SC pool and impairs muscle regeneration. A** Breeding scheme for generating inducible *Atf3* conditional knock out (*Atf3* cKO), *Pax7^Cre/+^; ROSA^EYFP/+^; Atf3^fl/fl^*, and control (Ctrl), *Pax7^Cre/+^; ROSA^EYFP/+^; Atf3+/+*, littermates. **B** Results of genotyping PCRs of the Ctrl and cKO mice. (F, *Atf3* Floxed; R, Cre recombinase in the *Pax7* locus; C, WT Ctrl). **C** The loss of ATF3 protein in the cKO FISCs was confirmed by western blot. α-Tubulin was used as a loading control. **D** Left: IF staining of Pax7 (red) and Laminin (green) at postnatal 28 and 56 days on uninjured TA muscles from the Ctrl or cKO mice. Scale bar: 50 μm. Right: Quantification of the numbers of Pax7+ SCs per 100 fibers; n = 3 mice per group. **E** Left: IF staining of Pax7 (green) and MyoD (red) on SCs from the Ctrl or cKO mice after cultured for 24 h. Scale bar: 25 μm. Right: Quantification of the percentage of Myod+/Pax7+ SCs; n = 3 mice per group.  **F** Left: SCs from Ctrl or cKO mice were cultured for 24 h before treating with EdU for 6 h and staining for EdU (red) and Pax7 (green). Scale bar: 50 μm. Right: Quantification of the percentage of EdU+/Pax7+ SCs; n = 3 mice per group. **G** H&E staining of the TA muscles collected at 5 and 7 dpi from Ctrl or cKO mice. Scale bar: 50 μm. **H** CSAs of newly formed fibers were quantified from the above-collected TA muscle at 7dpi and the distribution is shown; n = 3 mice per group. **I** Left: IF staining of eMyHC (red) and laminin (green) was performed on the above TA muscles collected at 5 and 7 dpi. Scale bar: 50 μm. Right: Quantification of the numbers of eMyHC+ fibers per view; n = 3 mice per group. All the bar graphs are presented as mean ± SD. ∗p < 0.05, ∗∗p < 0.01, ∗∗∗p < 0.001. n.s., no significance.

**Supplementary Figure 5.** **ATF3 deletion induces SC activation during voluntary and resistance exercises. A** Ctrl and iKO mice were subject to voluntary running wheels for 1 week and followed by 3 more weeks of running. The daily running distance of Ctrl and iKO mice were recorded. **B** The average running distance per day of Ctrl and iKO mice. **C** qRT-PCR detection of *Atf3, Atf4*, *Fos*, *FosB* and *Junb* in FISC isolated from Ctrl muscles -EX, +VE or +RE. All the bar graphs are presented as mean ± SD. ∗p < 0.05, ∗∗p < 0.01, ∗∗∗p < 0.001. n.s., no significance.

**Supplementary Figure 6. ATF3 regulates H2B gene expression and nucleosome patterning. A** RNA-Seq was performed in FISCs from the Ctrl and iKO cells and heat maps indicating differential gene expression levels(Log2[FPKM]) in the *Atf3* iKO vs. Ctrl. **B** RT–qPCR validation of the expression of selected histone genes in the iKO vs. Ctrl. **C** Illustration of the 2 clusters of Histone H2B coding genes on mouse chr3 and chr13. **D** RNA-Seq was performed in FISCs from the Ctrl and cKO mice. DEGs were identified in the cKO vs. Ctrl. **E** FPKM and Log2 fold change (FC) of *H2b* genes from the above RNA-Seq. **F&G** Gene ontology (GO) analyses of the above identified up and down-regulated genes in D. The top 10 enriched GO terms ranked by gene ratio (proportion of genes annotated for each GO term) are shown. Dots are colored by adjusted P value (fold change) and their size corresponds to the gene counts annotated to each GO term. **H** ATF3 was overexpressed in C2C12 with a pcDNA3.1-ATF3 plasmid. α-Tubulin was used as a loading control. **I** Genomic snapshots of 5 of the above identified *H2b* genes (*Hist1h2bf*, *Hist1h2bj*, *Hist1h2bk*, *Hist1h2l* and *Hist1h2bn*) with ATF3 binding in their TSSs (ChIP-Seq tracks) and down-regulated by ATF3 deletion (RNA-Seq tracks). **J** Left: Venn diagrams show the overlapping (112 genes) between the ATF3 ChIP-Seq target genes (2871) and the up-regulated genes (1866). Right: GO analysis of the above 112 genes. All the bar graphs are presented as mean ± SD. ∗p < 0.05, ∗∗p < 0.01, ∗∗∗p < 0.001.

**Supplementary Figure 7. Screening of other functional AP-1 family members in SCs and regeneration.** **A** Schematic outline of the experimental design for screening of functional AP-1 members. AAV9-sgRNA virus particles were intramuscularly injected into *Pax7^Cas9^* mice at P10. BaCl_2_ was injected to induce muscle regeneration 7 weeks later. TA muscles and FISCs were then collected at 5 dpi for staining and EdU assays respectively. **B-E** Upper: Illustration of locations of sgRNAs designed for targeting *Atf4*, *FosB*, *Fos* or *JunB* locus. Lower left: PCR analysis of the editing efficiency. WT(Wild-type fragments) and KD(Cas9 cleaved fragments) are indicated by arrows; n = 3 mice per group. Lower right: the protein levels of the above targeted AP-1 were examined by Western blotting in FISCs. GAPDH was used as a loading control. **F** The protein levels of ATF4, FOS, FOSB and JUNB in FISCs from Ctrl or KD mice were examined by IF staining. **G** The RNA levels of *Atf4*, *Fos*, *FosB* and *JunB* in the above FISCs were examined by RT–qPCR. **H** Left: SCs from *Ctrl* or *Atf4*, *FosB*, *Fos* and *JunB*-KD mice were cultured for 24 h. before treated with EdU for 6 h and stained for EdU (red) and Pax7 (green). Scale bar: 50 μm. Right: Quantification of the percentages of EdU+/Pax7+ SCs; n ≥ 3 mice per group. **I** Left: IF staining of Pax7 (red) and Laminin (green) on TA muscles from the above KD mice 56 days after AAV injection. Scale bar: 50 μm. Right: Quantification of the numbers of Pax7+ SCs per 100 fibers; n ≥ 3 mice per group. **J** H&E staining of the TA muscles collected from the above KO mice at 5 dpi. Scale bar: 50 μm. **K** Left: IF staining of eMyHC (red) and Laminin (green) was performed on the TA muscles from the above KD mice collected at 5 dpi. Scale bar: 50 μm. Right: Quantification of the numbers of MyHC+ fibers per view 5dpi; n ≥ 3 mice per group. **L** IF staining of Pax7 (red) and Laminin (green) was performed on the TA muscles from the KD mice collected at 5 dpi. Scale bar: 50 μm. Right: Quantification of the numbers of Pax7+ SCs per view at 5dpi; n ≥ 3 mice per group. **M** IF staining of MyoD (red) and Laminin (green) was performed on the TA muscles collected from the above KD mice at 5 dpi. Scale bar: 50 μm. Right: Quantification of the numbers of MyoD+ SCs per view at 5dpi; n ≥ 3 mice per group. All the bar graphs are presented as mean ± SD. ∗p < 0.05, ∗∗p < 0.01, ∗∗∗p < 0.001. n.s., no significance.
