## Supplementary material for "ATF3 Preserves Skeletal Muscle Stem Cell Quiescence by Preventing Precocious Activation": Suppl-figures

**Figure S1. Zhang S. et. al.**

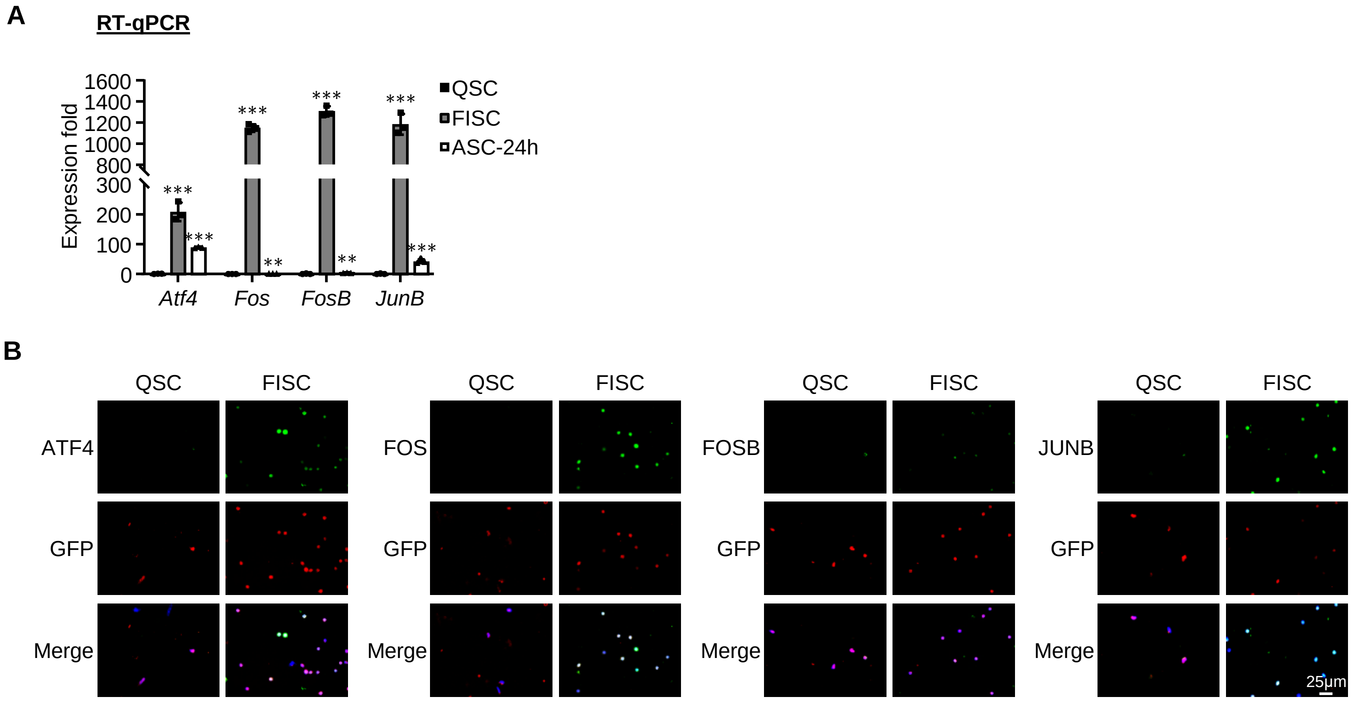

**Figure S2. Zhang S. et. al.**

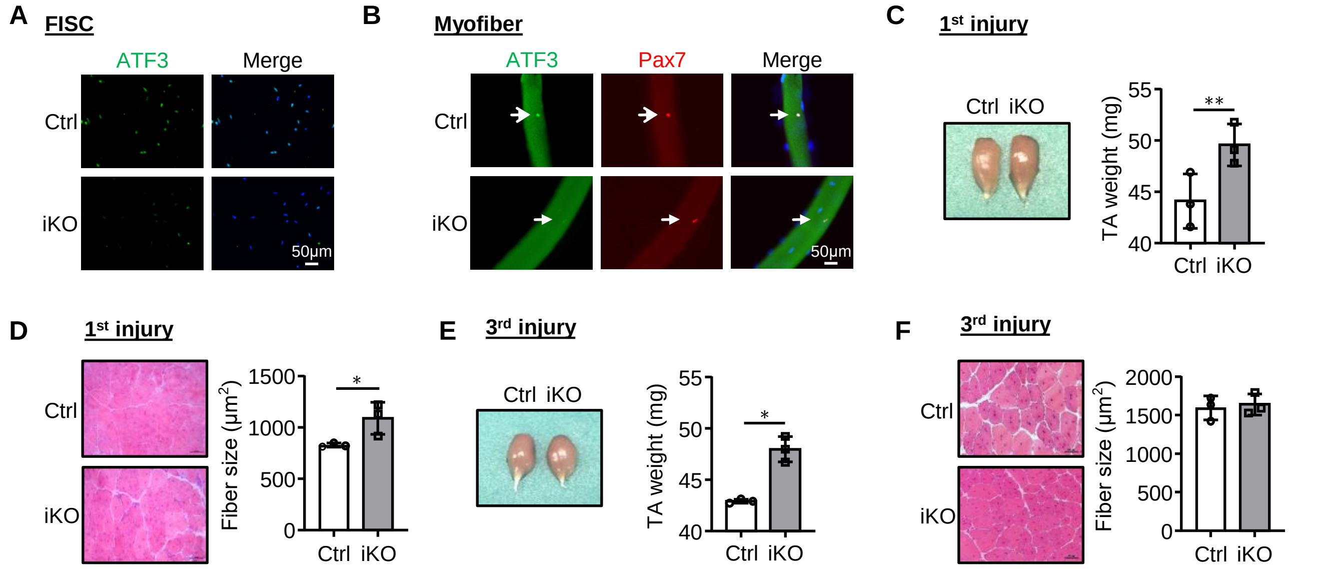

**Figure S3. Zhang S. et. al.**

**A** **SC-72h in culture**

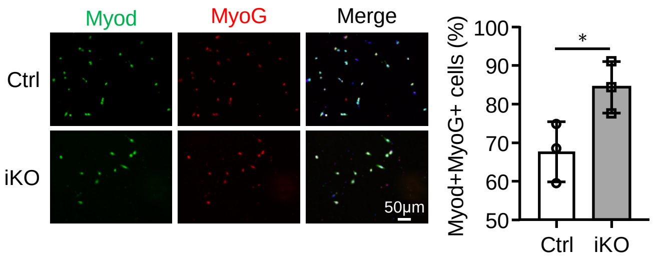

**Figure S4. Zhang S. et. al.**

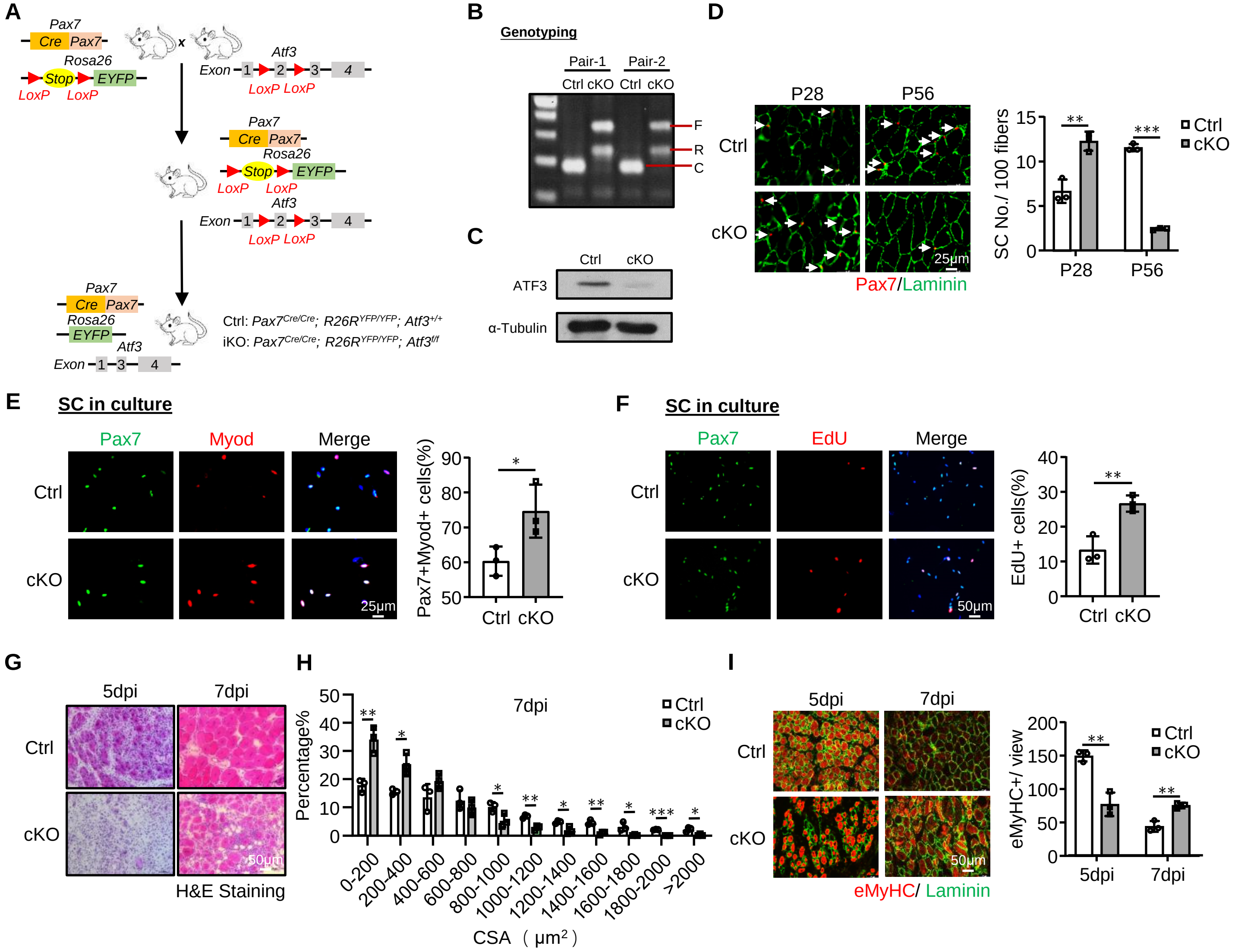

**Figure S5. Zhang S. et. al.**

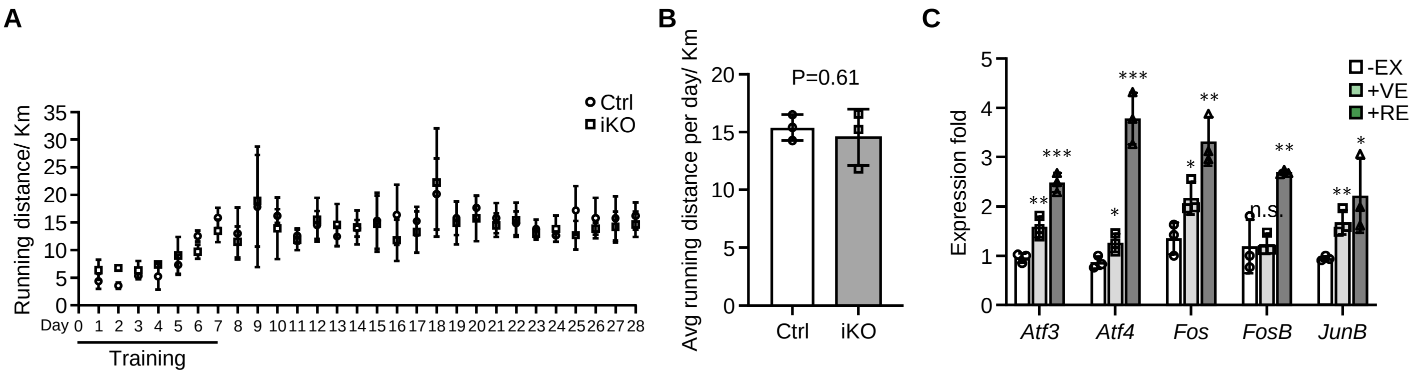

**Figure S6. Zhang S. et. al.**

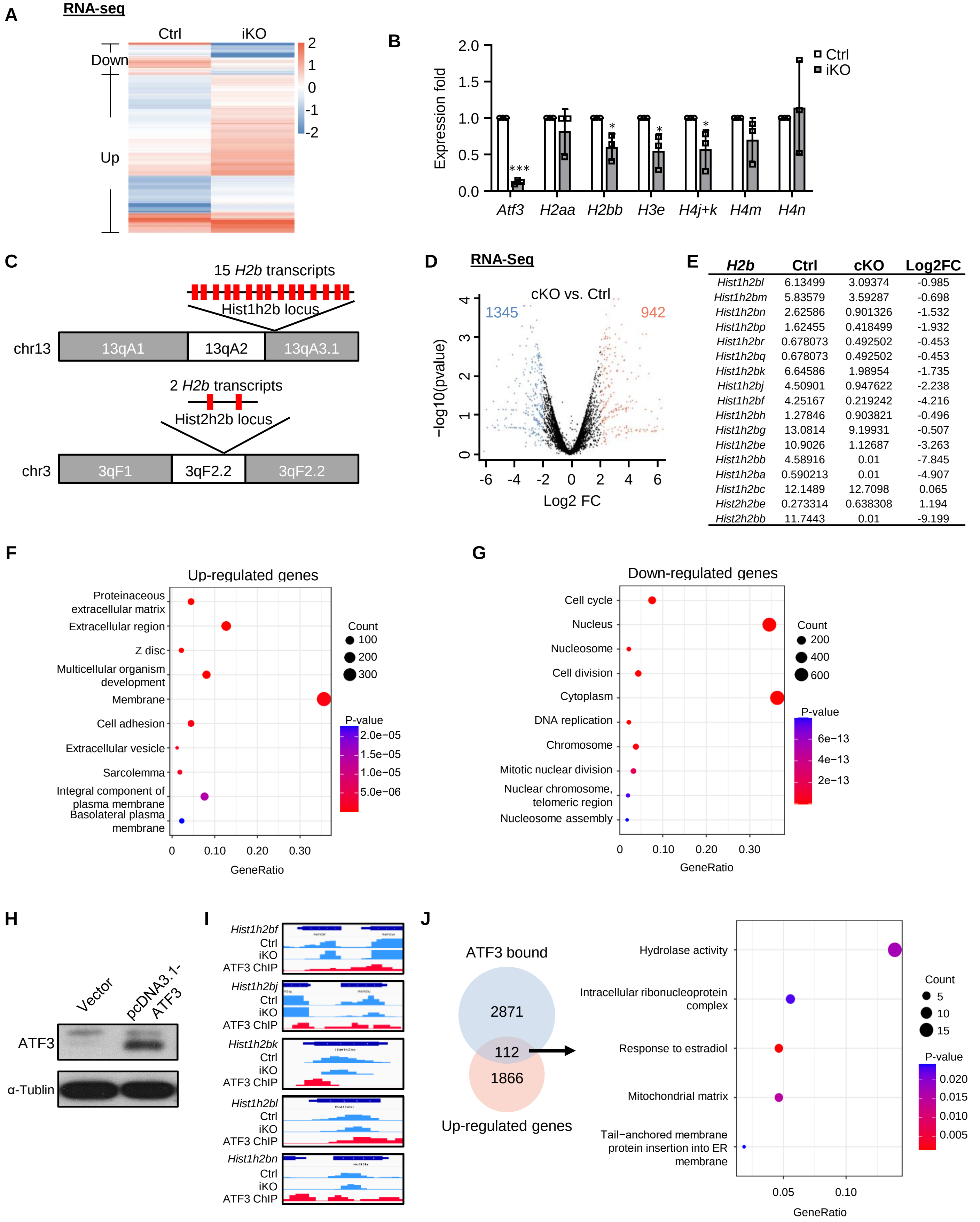


| *H2b* | Ctrl | cKO | Log2FC |
|---|---|---|---|
| *Hist1h2bl* | 6.13499 | 3.09374 | -0.985 |
| *Hist1h2bm* | 5.83579 | 3.59287 | -0.698 |
| *Hist1h2bn* | 2.62586 | 0.901326 | -1.532 |
| *Hist1h2bp* | 1.62455 | 0.418499 | -1.932 |
| *Hist1h2br* | 0.678073 | 0.492502 | -0.453 |
| *Hist1h2bq* | 0.678073 | 0.492502 | -0.453 |
| *Hist1h2bk* | 6.64586 | 1.98954 | -1.735 |
| *Hist1h2bj* | 4.50901 | 0.947622 | -2.238 |
| *Hist1h2bf* | 4.25167 | 0.219242 | -4.216 |
| *Hist1h2bh* | 1.27846 | 0.903821 | -0.496 |
| *Hist1h2bg* | 13.0814 | 9.19931 | -0.507 |
| *Hist1h2be* | 10.9026 | 1.12687 | -3.263 |
| *Hist1h2bb* | 4.58916 | 0.01 | -7.845 |
| *Hist1h2ba* | 0.590213 | 0.01 | -4.907 |
| *Hist1h2bc* | 12.1489 | 12.7098 | 0.065 |
| *Hist2h2be* | 0.273314 | 0.638308 | 1.194 |
| *Hist2h2bb* | 11.7443 | 0.01 | -9.199 |

**Figure S7. Zhang S. et. al.**

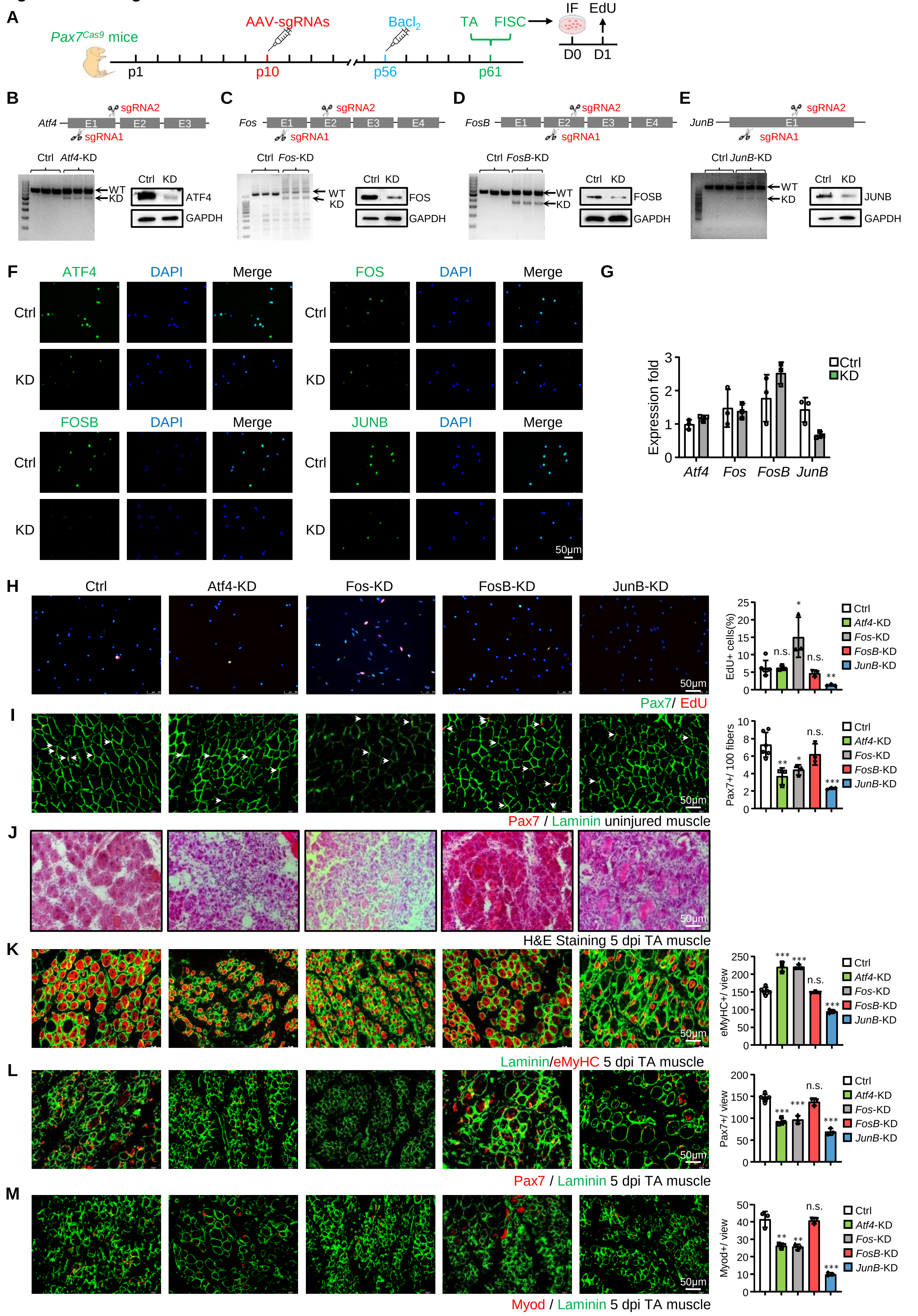
